# De novo discovery of structural motifs in RNA 3D structures through clustering

**DOI:** 10.1101/155580

**Authors:** Ping Ge, Shahidul Islam, Cuncong Zhong, Shaojie Zhang

## Abstract

As functional components in three-dimensional conformation of an RNA, the RNA structural motifs provide an easy way to associate the molecular architectures with their biological mechanisms. In the past years, many computational tools have been developed to search motif instances by using the existing knowledge of well-studied families. Recently, with the rapidly increasing number of resolved RNA 3D structures, there is an urgent need to discover novel motifs with the newly presented information. In this work, we classify all the loops in non-redundant RNA 3D structures to detect plausible RNA structural motif families by using a clustering pipeline. Compared with other clustering approaches, our method has two benefits: first, the underlying alignment algorithm is tolerant to the variations in 3D structures; second, sophisticated downstream analysis has been performed to ensure the clusters are valid and easily applied to further research. The final clustering results contain many interesting new variants of known motif families, such as GNAA tetraloop, kink-turn, sarcin-ricin, and T-loop. We have also discovered potential novel functional motifs conserved in ribosomal RNA, sgRNA, SRP RNA, riboswitch, and ribozyme.

## Introduction

Non-coding RNAs (ncRNAs) achieve their specific biological functions by folding into three-dimensional (3D) structures with many locally stable components. Among them, some highly abundant building blocks, called “RNA structural motifs”, are found to play important roles which may determine the behaviors of the molecules. For examples, the kink-turn motifs are the important binding sites for nine proteins in the bacterial 23S ribosomal RNAs (rRNAs);^1^ the cleavage at sarcin-ricin motifs led by the toxic proteins may result in complete shutdown of protein synthesis.^2^ Therefore, the identification and understanding of these recurrent structural components are indispensable for the study of RNA molecules. Considering that the number of resolved RNA 3D structures is rapidly increasing in recent years, thorough analysis of structural motifs is expected to extend our knowledge of the relationship between RNA structures and functions.

One major computational approach for studying RNA structural motifs is to search homologous instances of known motifs by using comparative methods. Traditionally, the motifs are modeled with their 3D geometric features, such as backbone conformations or torsion angles. NASSAM^3^ and PRIMOS^4^ are typical tools which primarily rely on the 3D atomic coordinates. They perform well for some simple motifs, but may not work for complex ones since the underlying computational methods are too rigid to identify the flexible variations in structures. Besides 3D information, FR3D integrates pairwise interactions as constraints into the screening for RNA structural motifs.^5^ However, as the most critical character of RNAs, the base-base interactions should be used as key factors in the assessment of structural discrepancy directly.^6^ Based on this idea, RNAMotifScan is proposed to search new motif candidates that share non-canonical base-base interaction patterns with the query.^7,8^ The benchmarking results show that RNAMotifScan outperforms other state-of-the-art RNA structural motif searching tools, especially for the instances with geometric variations caused by insertions or deletions.

An issue of searching tools is that they are based on the existing knowledge of RNA structural motifs, and thus cannot be used to detect new families. To tackle this problem, comparative methods are incorporated into clustering pipelines for the de *novo* discovery of conserved structural elements. One example is COMPADRES,^9^ which makes use of PRIMOS to categorize RNA structural motifs in the database of existing RNA 3D structures. Its performance is limited by the rigid comparison between loop regions, and the clustering results are hard to be applied to further research due to the complex models only covering 3D geometric information. LENCS (longest extensible non-canonical substructure) adopts a much simpler model which defines the RNA structural motifs as graphs of nucleotides interconnected by base pairs and glycosidic bonds.^10^ Thus the structural similarity of two motifs can be evaluated by the maximum isomorphic subgraph. With this measurement, a hierarchical clustering tree is built, and the motifs with similar base-pairing patterns are categorized by cutting it with a universal threshold. LENCS has successfully identified several putative new motifs in three rRNAs without using any tertiary information. But its sensitivity to potential structural variations is low, because only the same types of base pairs are allowed to be matched in the graphs. A recent approach of classifying RNA structural motifs takes into account all the hairpin and internal loops in the non-redundant RNA 3D structures.^11^ Based on FR3D, this pipeline aims at grouping the loop regions conserved in 3D space together with the help of pairwise interaction constraints. All the annotated motif instances and families are well organized in an online database named RNA 3D Motif Atlas.^11^ One issue of RNA 3D Motif Atlas is the rigid restriction on the 3D geometric discrepancy between cluster members. For example, the helix-end nucleotides of the query and target loops are required to be very close in 3D space after superimposition. As a result, it intends to categorize the highly similar components into numerous small groups, and may lose insights of the possible structural variations in a motif family. RNA Bricks is an RNA structural motif database which also stores the external molecular environment of RNA structural motifs, such as contacts with other RNA motifs, proteins, metal ions, and ligands.^12^ But its classification of RNA structural motif families still depends on the RMSDs between atoms, which restricts the capacity to identify potential variations. We also have developed a clustering framework named RNAMSC for *de novo/* RNA structural motif identification in rRNAs.^13^ To ensure the high coverage of base-pairing information on the RNA sequences, the base-pairing annotation of two different tools, MC-Annotate^14^ and RNAView,^15^ were combined. Then the non-canonical base pairs in the loops were compared according to their isostericity,^16^ and the statistically significant alignments were determined by using P-values which were inferred from the simulated background data. After that, the conserved candidate pairs with low P-values were summarized into a graph, in which the strongly connected subgraphs were retrieved. The experimental results show that RNAMSC not only outperforms LENCS in the recovery of known motifs, but also discovers several novel motif families. Compared with RNA 3D Motif Atlas, our approach adopts base-pairing information to measure structural similarity in the clustering. As a result, RNAMSC can detect potential motifs with higher structural flexibility.

Here we propose a new clustering pipeline to automatically detect novel RNA structural motifs by extending RNAMSC. First, the new pipeline is optimized for the large-scale inputs, such as the non-redundant RNA structure dataset. Second, all the single-strand regions in the RNA molecules, including the multi-way junctions, are considered in the classification. Third, the clustering results are post-processed to analyze their functionalities and relationships. By using this new clustering approach, we have identified 191 motif families, in which 68 are from hairpin loops, 77 are from internal loops, and 46 are from multi-loops. Generally, the large clusters contain the known motifs in RNA 3D structures, such as GNRA tetraloop, T-loop, kink-turn, and sarcin-ricin. The variations in some motif families which are accidentally separated from the majority can be retrieved based on checking their secondary and tertiary structural patterns in the downstream analysis. Furthermore, we also have discovered some novel motifs conserved in both rRNAs and non-rRNAs, such as single guide RNA (sgRNA) in Cas9 complex, Alu domain in the signal recognition particle RNA (SRP RNA), GlmS riboswitch, and twister ribozyme.

## 1 MATERIALS AND METHODS

### 1.1 Data preparation

Our clustering approach is designed to use the known knowledge of RNA 3D structures deposited in the PDB database.^17^ As the paper is written, there were over 3,000 experimentally resolved macro-molecular structures containing RNAs. To avoid possible biases in the statistical evaluation of structural conservation, the non-redundant (NR) list (of RNA-containing PDB structures) from the BGSU RNA group^18^ was adopted. This dataset eliminated the redundancy both in a single PDB file and among multiple PDB files, while keeping sufficiently diverged homologous structures. The selected 876 PDB files (including 1,307 RNA chains) at 4.0 *A* resolution threshold in v1.89 NR list were downloaded.

All the plausible pairing interactions in the RNA 3D structures were identified by using MC-Annotate^14^ and RNAView.^15^ Their annotation results were merged, and the conflicts were resolved by taking the MC-Annotation predictions. For each chain, the predicted *cis* Watson-Crick base pairs were retrieved to reveal the stacks in the RNA secondary structure. The pseudoknots in the structure were recognized by the program K2N^19^ and then eliminated. In the pseudoknot-free secondary structure, the single-strand regions were decomposed into hairpin loops, internal loops (including bulges), and multi-loops by the consecutively nested cis Watson-Crick base pairs (≥ 2). The loops without base-pairing interaction were removed to refine the datasets. Given the fact that some known structural motifs were closed by cis Watson-Crick base pairs, the helix ends were retained in the loops. Finally, the orders of the strands in loops were considered to generate motif candidates. The two strands in the internal loops were concatenated in both ascending and descending orders. For loops with more than two strands, we reduced the excessive number of permutations by converting them into circular forms in the 5’-3’ direction, and only the permutation in the cyclic orders were used in the furthur computation.

### 1.2 Loop alignment and clustering

All the motif candidates were grouped into three different datasets: HL (from hairpin loops), IL (from internal loops and bulges), and ML (from multi-loops). The HL dataset contained 1,036 instances, the IL dataset contained 1,868 instances, and the ML dataset contained 2,778 instances. In each dataset, an all-to-all alignment was performed by using RNAMotifScan. Because RNAMotifScan is developed for RNA structural motif searching, it treats queries and targets differently in the computation.

Thus, for any two candidates, one was aligned twice to its partner, as the query in the first alignment and as the target in the second alignment. The two corresponding Z-scores were computed with the alignment score distributions of queries, and the smaller one was assigned to the candidate pair as the numerical measurement of their tertiary structural similarity.

After that, three weighted graphs were constructed from the alignment results for different datasets. In these graphs, the vertices were the loops in RNAs and the edges were labeled with the alignment Z-scores. Note that internal loops and multi-loops have multiple candidates with different orientations of strands. The maximum Z-score of all candidate alignments for two loops was chosen as the weight of the edge. A Z-score cutoff was set to determine whether an edge should be removed or not. The remaining edges represent the highly significant structural conservation between loops, and the strongly connected sub-graphs were identified with a CAST-like clique finding algorithm.^20^

During the loop alignment and the clustering, the parameters of the pipeline were tuned to generate the most reliable results for motif discovery. For RNAMotifScan, thirty sets of parameters were used: the weights for sequence similarity and structural similarity can be (0.2, 0.8) or (0.4, 0.6); the gap start and extend penalties can be (3, 2), (6, 2) or (6, 4); the penalty of missing one base pair in two inputs can be 1 to 5. Different Z-score cutoffs, ranging from 1.0 to 3.0 with a step of 0.1, were also applied in the graphs. Therefore, there were 630 (30 x 21) different clustering results for each loop dataset (HL, IL, or ML). For HL and IL datasets respectively, we used the known motifs in rRNAs (1S72 and 1J5E)^13^ as a benchmark to select one clustering result with the highest specificity and sensitivity. The selected clustering result for HL uses following parameters: 0.2, 0.8, 6, 4, 3, and -1.1. The selected clustering result for IL uses following parameters: 0.2, 0.8, 6, 2, 2, and -2.2. For the ML dataset, since there is no enough known motif instances in multi-loops to conduct a solid analysis, the selected clustering result for ML used the same parameters selected for IL dataset.

### 1.3 Motif family identification

We extracted the potentially conserved motif families from the clusters for HL, IL, and ML datasets by using both computational methods and visual inspections. The clusters with size less than 3 were not considered in the further analysis. Each edge in a cluster must satisfy two requirements to be retained in the graph: First, the length of the loop region in the consensus structure derived from the alignment must be greater than 2. Second, the corresponding root-mean-square deviation (RMSD) must be less than 4A. After prouring all the invalid edges in the graph, the isolated loops in the cluster were removed. To distinguish the original clusters and the processed clusters, henceforth we will use “clusters” and “refined clusters” to refer to them, respectively. Furthermore, different motifs may be classified together if they share common patterns in their secondary structures.^8^ The 3D structures of the remaining loops in refined clusters were manually checked to categorize them into sub-clusters. After that, the secondary and tertiary structural features of sub-clusters were extracted for function annotation, and the identical sub-clusters were investigated through comparing these features. Finally, we designed an ID system to refer to the clusters and sub-clusters. The cluster ID contains two fields: a loop type prefix and a cluster index suffix (e.g. IL1). Based on that, the sub-cluster ID is defined as cluster ID followed by the sub-cluster index, separated by an underscore character (e.g. IL1_1).

## 2 RESULTS

### 2.1 Summary of the clustering results

We have identified 191 clusters whose sizes are greater than 2, among them 68 clusters from the HL dataset containing 600 loops, 77 clusters from the IL dataset containing 727 loops, and 46 clusters containing 203 loops. All the clusters and their members are listed in Supplementary Table S1. After removing the false conservations, the results have 57 refined HL clusters containing 473 loops, 57 refined IL clusters containing 463 loops, and 42 refined ML clusters containing 165 loops. Some of the refined clusters were further divided into sub-clusters. There are 68, 91, and 46 sub-clusters for the HL, IL, and ML datasets, respectively. Among them, 13 HL sub-clusters and 37 IL sub-clusters were annotated as known motif families. All the other sub-clusters, 55 from HL, 54 from IL, and 46 from ML, potentially belong to novel motif families. All the sub-clusters and the corresponding annotation information are listed in Supplementary Table S2.

To evaluate the performance of the automatic pipeline, 10 well-studied motif families in the clustering results were analyzed. The clusters containing the maximum numbers of instances for these motifs were chosen as representatives. All the other motif instances not in the representatives were annotated as plausible variations. Table 1 summarizes the benchmarking results for the 10 motif families, including the prediction accuracy (based on motif instances in the representatives) and the numbers of variations (based on motif instances not in the representatives). The precision was computed by dividing the number of true motifs with the size of the representative cluster. If one loop consists of several different motifs, it will be counted multiple times, as a true instance in representative and as variances of other motifs. From Table 1, we can see that the clustering results of GNAA and GNGA motifs are very accurate. This is because they are highly conserved in both sequences and secondary structures. The related variations mainly come from the hairpin loops with multiple motifs, which causes them to be categorized into other representatives. T-loops are relatively hard to be clustered together, due to the low sequence identity and simple base-pairing patterns. It indicates that the structural similarity weight should be set much greater than sequence similarity weight when searching T-loops in known RNA 3D structures. Both sarcin-ricin and kink-turn motifs have numerous variations, which are not classified with the majority of instances. One possible reason is that the binding activity may disturb the base-pairing interactions in them, and we will show several examples in the later sections. In addition, the precision of the kink-turn cluster (IL5) is relatively low because it contains several E-loops, whose secondary structure consensus is partially similar to the kink-turn’s.^7^ Hook-turn has a unique base-pairing pattern, so it is relatively easy to identify. Although C-loop is hard to detect due to the crossing base pairs, our pipeline still achieves acceptable results for it. All the other three motifs, E-loop, tandem shear, and reverse kink-turn, consist of continuous non-canonical base pairs. E-loop and tandem shear have similar 3D structures, so we mainly use their secondary structural features to distinguish them. Note that the previous accuracy analysis is based on the original clustering results. After the post-processing, the precisions of the sub-clusters for the benchmarking motifs are all 100%.

**Table 1.**
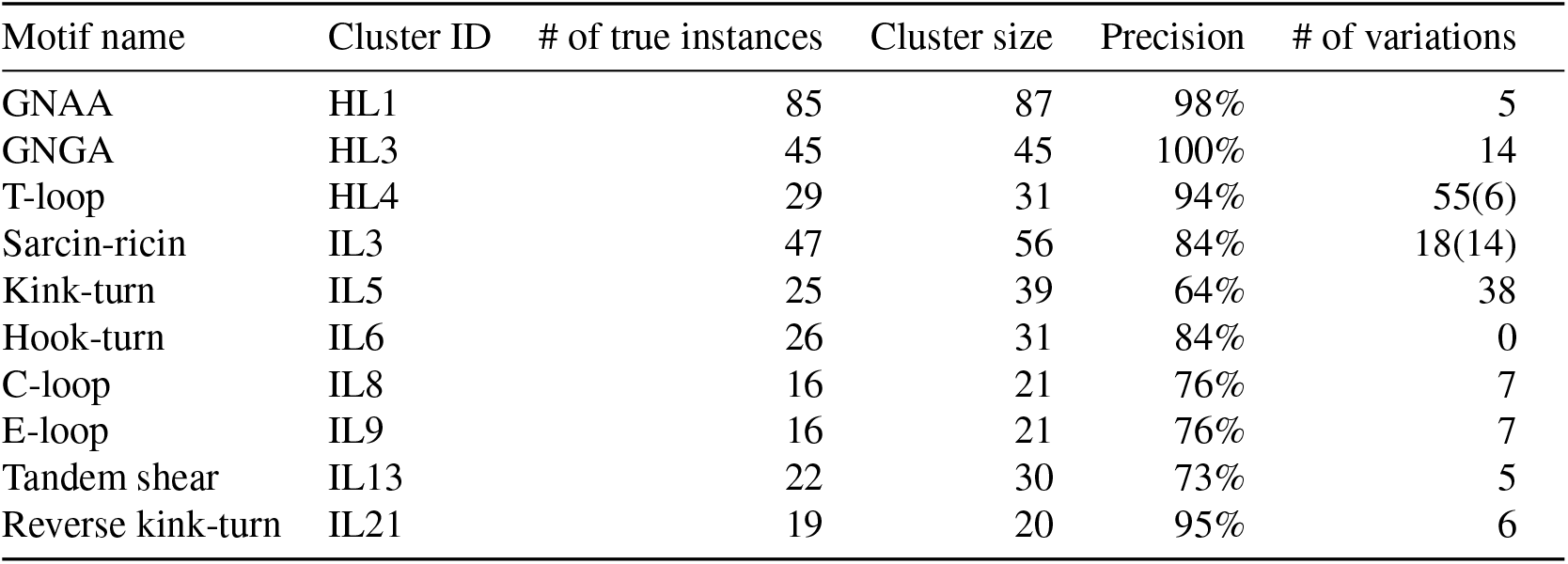
The clustering results of 10 well-known motif families.

| Motif name | Cluster ID | # of true instances | Cluster size | Precision | # of variations |
| --- | --- | --- | --- | --- | --- |
| GNAA | HL1 | 85 | 87 | 98% | 5 |
| GNGA | HL3 | 45 | 45 | 100% | 14 |
| T-loop | HL4 | 29 | 31 | 94% | 55(6) |
| Sarcin-ricin | IL3 | 47 | 56 | 84% | 18(14) |
| Kink-turn | IL5 | 25 | 39 | 64% | 38 |
| Hook-turn | IL6 | 26 | 31 | 84% | 0 |
| C-loop | IL8 | 16 | 21 | 76% | 7 |
| E-loop | IL9 | 16 | 21 | 76% | 7 |
| Tandem shear | IL13 | 22 | 30 | 73% | 5 |
| Reverse kink-turn | IL21 | 19 | 20 | 95% | 6 |

Besides the motifs in the table, we have discovered other functional ones in the clustering results. The first example is the well-known tetraloop receptor in group I intron (IL4_1 and IL22_1).^21^ Some of them are used in the target molecules to maximize their crystallizability.^22^ The L1 protuberance of 50S rRNA and mRNA were also clustered together in IL18. It has already been proved that they have both similar 3D structures and binding activities.^23^ We have also detected the motifs that are conserved both in mitochondrial 16S rRNAs and bacterial 23S rRNAs.^24^ The identification of these known functional motifs indicates that the clustering results can be applied to further analysis for new motifs.

### 2.2 Novel instances of known motifs

#### 2.2.1 Tetraloop

Tetraloops are the basic building blocks of RNA 3D structures. They are very important for thermodynamic stability and binding activity of the molecules.^25,26^ The most frequent two types of tetraloops are GNRA loops^27^ and UUCG loops.^28^ GNRA loops can be further categorized into GNGA loops and GNAA loops. In our clustering results, the majority of GNAA, GNGA, and UUCG motifs are in HL1, HL3 and HL6. Some other GNAAs and GNGAs co-exist with sarcin-ricin motifs in the loops. One instance of this motif module is shown in Figure 1 (a). This loop is from the region C3120-A3136 in the *Homo sapiens* mitochondrial 16S rRNA. It can be seen that the 3D structure of A3125-G3131 is highly conserved to a GNAA reference. We also found that the corresponding region in the *Haloarcula marismortui* 23S rRNA contains a GNGA motif (see HL35). Similar modules for UUCG have been also detected. One example is shown in Figure 1 (b), which is in the *H. marismortui* 23S rRNA. The “U-shape” turn in this loop is docked with the blue UUCG tetraloop precisely in the 3D space. Base on the observation, we may hypothesize that the combination of sarcin-ricin and tetraloop may be a very common module in RNA 3D structures.

**Figure 1.**
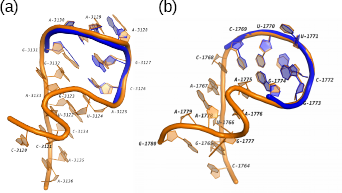
The 3D structures of two RNA motifs containing both tetraloops and sarcin-ricins. (a) The hairpin loop in the H. *sapiens* mitochondrial 16S rRNA (PDB: 3J7Y, chain: A) which contains a GNAA tetraloop. The blue tube shows a superimposed GNAA tetraloop in a 23S rRNA (PDB: 4BW0, chain: A, 9-14). (b) The hairpin loop in the H. *marismortui* 23S rRNA (PDB: 1S72, chain: 0) which contains a UUCG tetraloop. The blue tube shows a superimposed UUCG tetraloop in a mRNA fragment (PDB: 2HW8, chain: B, 15-20).

#### 2.2.2 T-loop

T-loop is a compact U-turn-like loop which was originally discovered in tRNA.^29^ After that, many T-loop instances have been identified in a variety of ncRNAs, ranging from rRNA to riboswitch.^30^ Our clustering results cover almost all the known T-loops in the hairpin loops. What’s more, we also found two new instances of T-loop in the internal loops (IL26). One of them is in the Thi-box (thiamine pyrophosphate sensing) riboswitch and known for the ligand 4-amino-5-hydroxymethyl-2-methylpyrimidine (HMP).^31^ The other one is in a T-box stem I RNA. Figure 2 shows its structure and the corresponding 3D docking to a T-loop in a tRNA. Note that both secondary structures consist of one *trans* S/H and one *trans* W/H base-pairing interactions. The difference is that in the tRNA the base pairs exist in a hairpin loop, while the two interactions in the T-box stem I RNA bend one strand of the internal loop to a U-shape turn. Considering the relatively large size of the twisted strand, the third interaction at G38 homology is 4MGN_C:51-63 in HL8) in the same RNA to stack on tRNA elbow.^32^ The similar binding behavior is also found in RNase P and rRNA, so the study of this T-loop and its partner may provide useful information for searching more functional modules.

**Figure 2.**
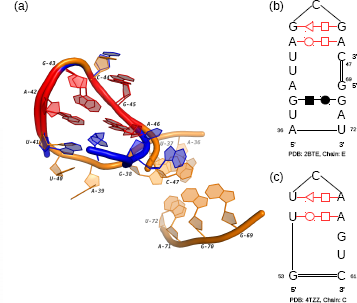
The 3D and secondary structures of an internal loop in T-box stem I RNA and a hairpin loop in tRNA. (a) The 3D docking of two loops. The orange tube represents the internal loop in the T-box stem I RNA (PDB: 4TZZ, chain: C) and the blue tube represents the hairpin loop in the tRNA (PDB: 2BTE, chain: E). (b) The secondary structure of the internal loop. (c) The secondary structure of the hairpin loop. In (a), (b), and (c), the conserved base-pairing interactions are marked in red.

#### 2.2.3 Kink-turn

Kink-turn is a motif in the internal loop region with an asymmetrical architecture.^1^ Its key feature is the tight kink at the backbone of the longer strand, which causes the axes of the two helical stems differ by about 120 °. In a real cellular environment, kink-turn may adopt a dynamic conformation.^33^ To maintain the specific 3D geometry, the motifs require the presence of metal ions,^34^ or binding with proteins.^35^ We detected two new kink-turn-like motif instances in the cluster IL37. Note that the loop in the 16S rRNA was detected in our previous work.^13^ With the newly discovered instance, we can analyze their conserved patterns and the related functions. Figure 3 shows their secondary and 3D structures. It can be seen that all base pairs can be matched. Compared with the base-pairing pattern of common kink-turns,^8^ these two instances form the kinks by three base pairs (G247/A282, A246/G278, A246/G281 in 1FJG and G18/A48, A17/A44, A17/G47 in 3RW6). In the common kink-turns, the Watson-Crick base pairs, C242/G284 in 1FJG and U11/G50 in 3RW6, should be followed by two continuous non-Watson-Crick base pairs. However, in these two loops, the two interactions are separated by the nucleotides marked with red color. In Figure 3, these red nucleotides form the bulges in the shorter strand, which do not exist in the common kink-turns. What’s more, both red regions have long-range interactions. According to the results of MC-Annotate, the nucleotide U244 in 1FJG is paired with A893 in another loop region. On the other hand, the large bulge in 3RW6 containing the flipped out nucleotides, A13, G14, and A15, is the binding site of the TAP protein and critical to the formation of CTE-TAP complex.^36^ Based on the function similarity, we suggest that the secondary structural pattern is important for the long-range interactions of the kink-turn motifs.

**Figure 3.**
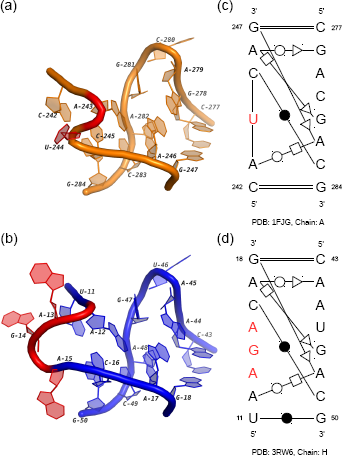
The 3D and secondary structures of two internal loops in a 16S rRNA and a CTE rRNA. (a) and (b) are the 3D structures of two loops in the 16S rRNA (PDB: 1FJG, chain: A) and the CTE RNA (PDB: 3RW6, chain: H). (c) and (d) are the secondary structures of (a) and (b). In (a), (b), (c), and (d), the nucleotides with binding activities are marked in red.

#### 2.2.4 Sarcin-ricin

Sarcin-ricin motif is first found in the large ribosomal subunit as the attacking site of two protein toxins, ricin and a-sarcin. The catalyzation among them will impact the binding between elongation factors and ribosome, which may result in the cessation of the protein synthesis.^37^ More sarcin-ricin instances with similar structural features have been discovered in other RNAs, including 5S and 16S rRNAs, by using computational methods.^7,38^ In our clustering results, the majority of sarcin-ricins were also detected in rRNAs (see the cluster IL3). Their secondary structures are almost the same as the widely used consensus,^38^ and the 3D structures are highly conserved with the known instances. On the other hand, we have also found some new functional loops that share structural features with sarcin-ricin. Here, we present two possible variations of sarcin-ricin whose secondary and 3D structures are shown in Figure 4. The first loop is in the cluster IL38. Based on the secondary structure, the S-shape turn in its 3D structure is mainly supported by two non-canonical base pairs (A415/G428 and A414/A430) and one outward stacking interaction (G428/A430). All these three pairing interactions are in the secondary structural consensus of sarcin-ricins.^8^ However, in common sarcin-ricins, the cis H/W base pair U429/A431 should be a cis H/S interaction between U429/A430. The possible reason for this difference is that the strand U427→C433 is longer than it in the consensus. What’s more, this motif instance contains a bulge at the strand G409→G416, which interacts with the S4 protein at G410, A411, and A412.^39^ So the two base-pairing interactions not in the consensus, A411/A430 and G413/G428, may be important for the maintaining of the long-range linkages. We may hypothesize that this motif is a sarcin-ricin variation whose secondary and 3D structures are disturbed by the protein binding activity. And the comparison of its secondary structure pattern with the sarcin-ricin consensus may help us to detect potential RNA-protein interactions.

**Figure 4.**
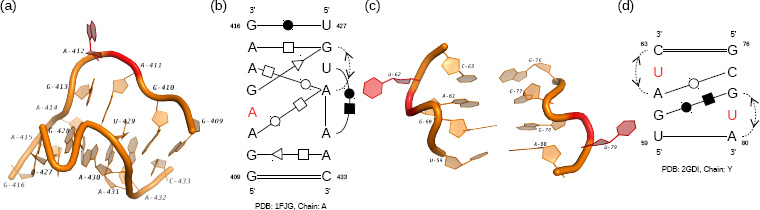
The 3D and secondary structures of two internal loops in a 16S rRNA and a TPP riboswitch. (a) The 3D structure of the internal loop in the 16S rRNA (PDB: 1FJG, chain: A). (b) The secondary structure of the loop in (a). (c) The 3D structure of the internal loop in the TPP riboswitch (PDB: 2GDI, chain Y). (d) The secondary structure of the loop in (c). In (a), (b), (c), and (d), the nucleotides with binding activities are marked in red.

Another interesting loop is in the cluster IL62. We call it “double S-turns” because there are two symmetrical S-shape turns in its 3D structure (see Figure 4 (c)). In the existing model for ligand-induced folding of the TPP riboswitch, this loop is the TPP-bind pocket which is critical for the ligand recognition.^31^ The two nucleotides, U62 and U79, shape the pocket by protruding into solution and weakening the stacking effects to the adjacent bases. From Figure 4 (d), it can be seen that there are two stacking interactions, A61/C63 and G78/A80, to enforce the local stability around these two nucleotides. They also cause the large turns in the S-shape structures. On the other hand, the other two non-canonical base pairs tighten the two strands together. The analysis of this internal loop, as well as the motif in Figure 4 (b), indicates that the stacking effect between discontinuous bases is an important evidence of detecting specific structural motifs, such as bulge and S-turn. In addition, this specific organization of interactions, including pairing interactions and stacking interactions, may be important to form pocket-like 3D structures.

### 2.3 Novel motif families

#### 2.3.1 Novel motif families in the hairpin loop regions

The first potential motif family mainly contains four different instances from HL2_1 and HL53_1. One of them is the loop 10 in yeast 18S rRNA.^40^ The other three are the “stem loop 1” in the sgRNA of the Cas9-sgRNA-DNA ternary complex.^41,42^ The mutations of residues interacting with stem loop 1 result in decreased DNA cleavage activity of the CRISPR-Cas system, which indicates the loop is essential for the formation of the functional Cas9-sgRNA complex. Figure 5 shows the high similarity between these two internal loops in terms of both geometric and base-pairing patterns. Except C275/G281 in 3U5F and G54/C60 in 4OO8, all the other interacted bases are identical in two loops. The continuity of the stacks is broken by U280 and U59 (labeled red color in Figure 5). Both of them flip out from the stems and cause the turns in the backbone of two loops. The most important feature is that they have similar functional roles: U280 interacts with L24e protein through the eB13 bridge in the hyper-rotated state;^43,44^ U59 in the sgRNA hydrogen bonds with Asn77 in the bridge helix of Cas9.^42^ So, these loops are not only conserved in 3D structures but also in the functions, which implies the potential closely relationship between the base-pairing pattern and the protein binding activity.

**Figure 5.**
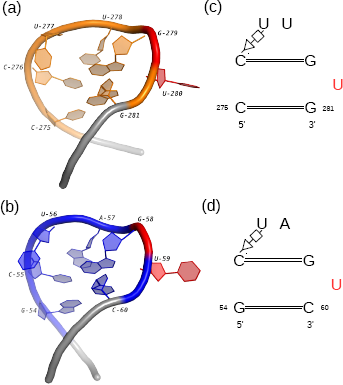
The 3D and secondary structures of two internal loops in a 18S rRNA and a Cas9-sgRNA-DNA complex. (a) and (b) are the 3D structures of two loops in the 18S rRNA (PDB: 3U5F, chain: 6) and the sgRNA (PDB: 4OO8, chain: B). The extended regions are shown in gray. (c) and (d) are the secondary structures of (a) and (b). In (a), (b), (c), and (d), the nucleotides binding with the proteins are marked in red.

#### 2.3.2 Novel motif families in the internal loop regions

IL16_1 contains four conserved regions in the 16S rRNAs and two loop B in the 5S rRNAs. We choose two representatives and describe their 3D and secondary structures in Figure 6. From Figure 6 (a) and (b), we can see that the common base-pairing interactions in two loops, which are shown in red, are highly conserved in 3D space. The corresponding base pairs in the secondary structures are from the same groups in the isostericity matrices:^16^ U375-A389 and G56-C26 belong to *cis* W/W I_1_; A374/C390 and A55/A27 belong to *trans* W/S I_1_; A373/G371 and A54/U52 belong to *trans* H/S I_1_. Therefore, they are co-varying mutations, and the interchange between them will maintain the 3D structures of the loops. What’s more, although the interactions of C372/A389 *(trans* W/H) and U53/C26 *(cis* H/W) are not from the same isosteric group, the geometric relationship of bases in them are quite similar.^45^ Therefore, these two base pairs may also contribute to the 3D structural similarity of these two internal loops.

**Figure 6.**
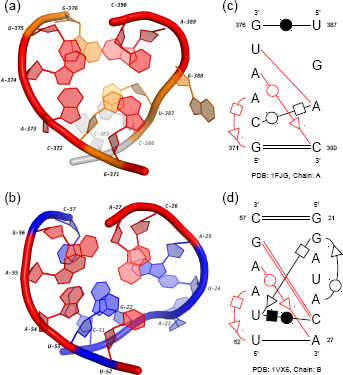
The 3D and secondary structures of two internal loops in a 16S rRNA and a 5S rRNA. (a) and (b) are the 3D structures of two loops in the 16S rRNA (PDB: 1FJG, chain: A) and the 5S rRNA (PDB: 1VX6, chain: B). The extended region in (a) is marked in gray. (c) and (d) are the secondary structures of (a) and (b). In (a), (b), (c), and (d), the conserved base-pairing interactions are marked in red.

The major difference between two motif instances comes from the regions U387-G388 and G21-A25. First, the lengths of two regions are different, which suggests a potential insertion in the loop of 5S rRNA. A significant feature shared between them is the turn on the phosphate backbone (Figure 6 (a) and (b)). However, the backbone of the internal loop in 5S rRNA (the blue one) turns with a slightly large angle. The reason may be the *trans* H/S base-pairing interaction between G22 and U53. Although the 3D structures of two regions are not totally the same, they actually may have similar molecular functions. Based on the results of MC-Annotate, the nucleotide G388 in the 16S rRNA, which flips out from the stem, interacts with C58. For the region in the 5S rRNA, a possible contact to helix 89 in 23S rRNA has been identified by a SELEX (systematic evolution of ligands by exponential enrichment) experiment.^46^ It is hypothesized that A23 is the possible binding site due to its base twisting further than the backbone. Moreover, the base-pairing consensus we detected here may be very critical for their interlinking functions.

Another possible novel functional motif is discovered in the cluster IL42. One instance in this cluster is from the 16S rRNA of *Thermus thermophilus,* while the other two are identical internal loops in the GlmS riboswitch of *Bacillus anthracis.* Riboswitches are metabolite-sensing RNAs that can directly control the expression of downstream genes.^47^ By binding to specific ligands, their structures are changed to terminate the transcription or hinder the translation. However, unlike other riboswitches, the GlmS riboswitch does not alternate its structure upon the binding of glucosamine-6-phosphate (GlcN6P).^48^ Instead, the binding activity results in a cleavage on the GlmS mRNA which reduces the GlcN6P synthetase production greatly.^49^ So it is also called “GlmS ribozyme”. The internal loop studied here interlinks two helices, P4 and P4.1, in the GlmS riboswitch. Its secondary and 3D structures are aligned with the loop in 16S rRNA, and the results are shown in Figure 7. We can see that although both strands of the loop in Figure 7 (a) are shorter than those of the loop in Figure 7 (b), the consensus marked by red color is highly conserved in sequences, base-pairing interactions, and 3D structures. The “S-shape” turns in the regions C1284-A1287 and U96-A98 are important common features of two loops too. In the GlmS riboswitch, the turn is supposed to pack obliquely into the minor groove of P2.1 helix, which is important for the GlcN6P binding.^50^ On the other hand, we also find that the flipped out nucleotide A1287 in the 16S rRNA also forms two interactions with A1353 and A1370. So the bulge-like structures may be indicators for the long-range tertiary interactions. The discovered motif may also be critical for the stability of the large internal loops whose structures are disturbed by intra-molecular linkages.

**Figure 7.**
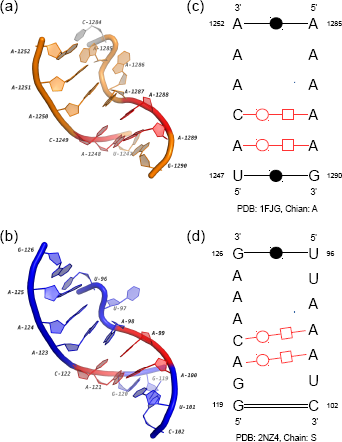
The 3D and secondary structures of two internal loops in a 16S rRNA and a GlmS riboswitch. (a) and (b) are the 3D structures of two loops in the 16S rRNA (PDB: 1FJG, chain: A) and the GlmS riboswitch (PDB: 2NZ4, Chain: S). The extended region in (a) is marked in gray. (c) and (d) are the secondary structures of (a) and (b). In (a), (b), (c), and (d), the conserved base-pairing interactions are marked in red.

#### 2.3.3 Novel motif families in the multi-loop regions

The first potential novel family in multi-loops is obtained from the sub-cluster ML2_1. Ten members are the orthologous regions from 21S, 23S, 25S, and 28S rRNAs, and the last one comes from the Alu domain of a signal recognition particle (SRP) RNA *(Bacillus subtilis).* SRP is a highly diverse ribonucleoprotein complex existing in all three kingdoms of life.^51^ The RNA in it can be divided into two functional domains, and one of them, the Alu domain, arrests protein biosynthesis by blocking the elongation factor entry site.^52,53^ Then by hindering the translation, SRP can prevent membrane proteins from being prematurely released from the ribosome. The multi-loop in the cluster is the one interlinking helix 1, helix 2 and helix 5a in the SRP RNA. Figure 8 shows the comparison of its secondary and 3D structures with the loop in 23S rRNA. Both loops have three strands, which are inter-connected by three highly conserved non-canonical base pairs: G1681/A1414 *(trans* S/H), A1414/A1682 *(trans* W/W) and A1682/U1696 *(trans* H/W) in 1S72, G62/A12 *(trans* S/H), A12/A64 *(trans* W/W) and A64/U101 *(trans* H/W) in 4WFL. The eight interacted nucleotides are marked with red color in Figure 8 (a) and (b). Note G62 and A64 in 4WFL. The structural difference between the Alu domain of SRP RNA in mammalian and bacteria may explain the potential function of the motif. The G-A-A-U four-base platform observed in bacteria *(B. subtilis)* is absent from the Alu domain in eukaryota. Previous experiments have already shown that the 5’ region of human Alu domain is very flexible and SRP9/14 proteins are required to stabilize the conformation and induce the binding to 50S rRNA.^54^ On the other hand, the bacterial Alu domain adopts a closed conformation directly with the help of the four-base platform. This evidence may suggest that the discovered motif is critical to the stabilization of the local structure that binds to proteins.

**Figure 8.**
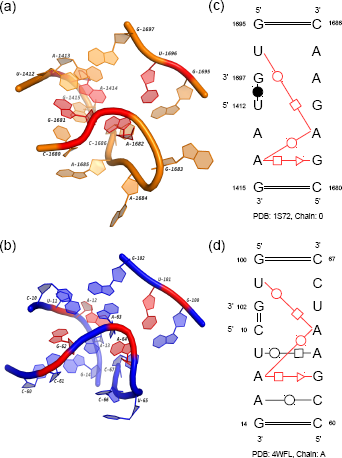
The 3D and secondary structures of two multi-loops in a 23S rRNA and the Alu domain of a SRP RNA. (a) and (b) are the 3D structures of two loops in the 23S rRNA (PDB: 1S72, chain: 0) and the SRP RNA (PDB: 4WFL, chain: A). (c) and (d) are the secondary structures of (a) and (b). In (a), (b), (c), and (d), the conserved base-pairing interactions are marked in red.

ML17_1 contains three conserved regions in 23S rRNAs and one instance in *env22* (type P1) twister ribozyme. As a small self-cleaving ribozyme, twister presents in many species of bacteria and eukaryota.^55^ Further research shows that twister may play a similar role as the hammerhead ribozyme in the biological systems. The instances of twister are categorized into three groups, type P1, type P3, and type P5, which can circularly permute to each others. The crystal structure of the twister used here comes from a type 1 instance. To compare it with the multi-loop in 23S rRNAs, we picked the one in 1S72 as a representative. Figure 9 shows the secondary structures of two loops and the 3D superimposition of their extensions. One common feature is that both loops are linked by *trans* S/S base-pairing interactions (A1492/C1514 and A42/C14). What’s more, the neighbors of the paired bases (G1512 and C1513in 1S72, G12 and C13 in 4RGE) form pseudoknots with nucleotides outside of the multi-loops (C1450 and G1449 in 1S72, C37 and G36 in 4RGE). Their 3D structures are highly conserved (red in 9). Note that the yellow multi-loop in 1S72 has four single-strand regions, while the blue one only has three. The extension of one strand in the yellow loop (C1455→G1453) involves in the formation of the pseudoknot. Although not a direct substitution, the blue loop has one sharply bent strand (G25→A26) who makes an 180° turn to serve the interaction. This interesting case in which the loops with completely different secondary structures have highly similar 3D structures may suggest the underlying tertiary structural pattern is very important.

**Figure 9.**
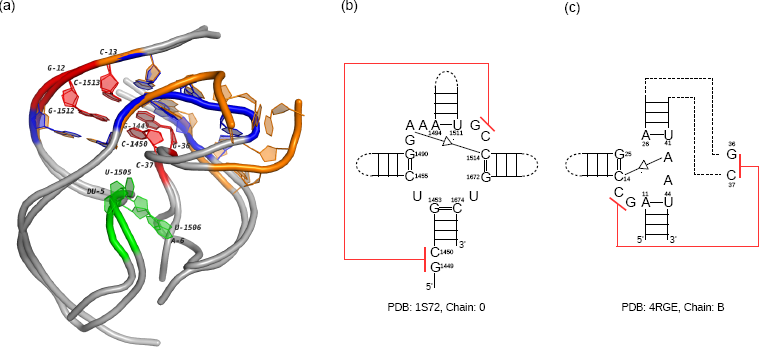
The 3D and secondary structures of two multi-loops in a 23S rRNA and an *env22* twister ribozyme. (a) The 3D docking of two loops. The yellow tube represents the multi-loop in the 23S rRNA (PDB: 1S72, chain: 0) and the blue tube represents the multi-loop in the twister ribozyme (PDB: 4RGE, chain: B). The extended regions of both loops are shown in gray. The nucleotides involved in the pseudoknots are labeled (G1512-C1450 and C1513-G1449 in 1S72, G12-C37 and C13-G36 in 4RGE). (b) The secondary structure of the multi-loop in the 23S rRNA. (c) The secondary structure of the multi-loop in the twister ribozyme. The pseudoknots in (b) and (c) are marked in red. The self-cleavage sites in 4RGE and the corresponding nucleotides in 1S72 are marked in green.

We also extend the 3D docking to the P2 and P4 helices of the twister ribozyme to study its local structural similarity with the 23S rRNA. Figure 9 (a) shows that the two RNAs are quite conserved in these 40-nt regions. The self-cleavage sites in the twister, dU5 and A6, are highlighted with green color. During the transcription, guanosine and Mg^2^+ are coordinated to the non-bridging phosphate oxygen at the U-A step for cleavage catalysis and structural integrity. The corresponding nucleotides, U1505 and U1506 in 1S72 (green) share a similar splayed-apart conformation with the cleavage sites in the twister ribozyme. With so many common features, these two regions should be further studied with the experimental effort to confirm their functional correlation.

## 3 Discussion and Conclusion

In this paper, we study the RNA structural motifs in non-redundant RNA 3D structures by using a *de novo* clustering approach. The single-strand regions in the corresponding secondary structures were extracted and categorized into hairpin loops, internal loops, and multi-loops. The base-pairing patterns in the same type of loops were compared by RNAMotifScan, and then the significant conservations were assembled into a graph. The densely connected sub-graphs were retrieved to form the clusters in which the members share common secondary structural features. In each cluster, by evaluating the alignments, the loops not close to any others in 3D space were removed. The remaining loops in the clusters were further analyzed, and then classified into different sub-clusters if their 3D structures were distinguishable from critical conformations. Finally, we tried to detect the homologous sub-clusters in different clusters by measuring the similarity of their secondary and 3D structural patterns. The clustering results for the known motifs indicate the high prediction accuracy of this new pipeline. Some interesting instances, which not only maintain the key features of known motifs but also exhibit specific structural variations, were found in the downstream analysis. We also identified numerous novel motif families, even in the multi-loop regions.

The in-depth investigation of the clusters provides directions for the further research. First, RNA structural motifs may work together as a “module”, such as the hairpin loops containing sarcin-ricins and tetraloops (see Figure 1), and the two T-loops in the T-box stem I RNA (see Figure 2). However, all the existing searching tools only focus on detecting the single motifs in isolation. Therefore, a new tool for discovering motif modules may provide essential evidence of the relationship among RNA structural motifs, which is important for the study of RNA structures and their functions. Another problem is to use base-pairing interactions to infer the potential binding activities between RNAs and other molecules. The disturbed secondary structures of the kink-turn and sarcin-ricin variations (see Figure 3 and Figure 4) reveal that they may be the indicators of the long-range linkages. Furthermore, the affected base pairs also have specific patterns which can be easily integrated into computational methods. This approach should be more accurate than the other methods based on indirect measurements, such as using the distances between atoms.

## 4 SUPPLEMENTARY DATA

Supplementary Tables S1 and S2.

